# NK cells mediate preventive efficacy of intravenous BCG against lung metastasis in mice

**DOI:** 10.1101/2024.10.09.617367

**Authors:** Claudia Guerrero, Eduardo Moreo, Miguel Araujo-Voces, Santiago Uranga, Ana Belén Gómez, Carlos Martín, Nacho Aguiló

**Affiliations:** Grupo de Genética de Micobacterias, Departamento de Microbiología y Medicina Preventiva, Facultad de Medicina, Universidad de Zaragoza, IIS-Aragon; Zaragoza, Spain; CIBER Enfermedades Respiratorias, Instituto de Salud Carlos III; Madrid, Spain; Departamento de Bioquímica y Biología Molecular, Instituto Universitario de Oncología (IUOPA), Universidad deOviedo, Oviedo, Spain

## Abstract

Lung metastases frequently arise from primary tumors, including bladder cancer, and represent a critical negative prognostic factor. Natural Killer (NK) cells have shown to play a vital role in controlling metastasis. Consequently, tumor cells have evolved specific mechanisms to evade NK cell-mediated immune surveillance, promoting metastasis and resistance to immunotherapy. In this study, we investigated the prophylactic and therapeutic potential of intravenous Bacillus Calmette–Guérin (BCG) in preventing lung metastases from bladder cancer cells using a murine model. We demonstrated that prophylactic BCG administration significantly reduced tumor burden and prolonged survival, largely through NK cell activation. However, BCG treatment was ineffective when administered over established tumors, likely due to tumor-driven immune evasion mechanisms. Our results revealed the contribution of interferon-gamma (IFN-γ) to tumor resistance. Tumor cells exposed to IFN-γ upregulated were more resistant to BCG *in vivo*, which correlated with the overexpression of immune checkpoint molecules, whereas disruption of the IFN-γ signaling pathway in tumor cells partially restored the therapeutic efficacy of BCG. Our findings highlight the importance of understanding tumor immune escape mechanisms and suggest that BCG could be a promising treatment for preventing lung metastases in bladder cancer.

## INTRODUCTION

Lung metastases commonly originate from a wide range of tumors, including bladder cancer^1^. Due to its extensive vascularization, the lung is particularly susceptible to the dissemination of cancer cells via the bloodstream or lymphatic system. As a result, the lungs are one of the most frequently affected organs in metastatic disease. Notably, the presence of lung metastases is a significant negative prognostic factor, leading to a substantial reduction in patient survival rates^2^.

Multiple clinical studies support the role of NK (Natural Killer) cells in the control of metastasis. Elevated expression of NK cell activation receptors or increased NK cell cytotoxicity has been correlated with favourable prognoses in several cohorts of cancer patients with metastatic disease or at high risk of developing it. For example, patients with primary prostate carcinomas whose infiltrating NK cells exhibit high levels of NKG2D and NKp46 have been shown to remain metastasis-free for at least one year after surgery^3^. Moreover, circulating NK cells have been identified as crucial sentinels in eliminating migrating tumor cells, thereby inhibiting the implantation of metastatic cells in distant organs^4^.

It has been described that tumor cells acquire mechanisms to specifically undermine NK cell activity, promoting cancer progression and the development of metastases. These mechanisms include the downregulation of ligands for NK cell activation receptors ^5^, along with the increased expression of specific immune checkpoints such as HLA-E and HLA-G ^6,7^ Additionally, tumor cells secrete regulatory cytokines, including TGF-β and IL-10, to diminish the cytotoxic function of NK cells.^8,9^, or the recruitment of immunosuppressive cell types such as regulatory T cells (Tregs) ^10^ and myeloid-derived suppressor cells (MDSCs) ^11^, which further inhibit NK cell functions. Moreover, hypoxia can influence the antimetastatic capacity of NK cells through the expression of hypoxia inducible factor (HIF-1)^12^ and the release of exosomes that contain TGF-β^13^ and activate autophagy^14^, which supress the activity of NK cells. Finally, it has been reported that metastatic tumor cells tend to migrate through circulation in clusters or are shielded by myeloid suppressor cells, allowing them to evade the cytotoxic action of NK cells in circulation ^15^.

BCG (Bacillus Calmette–Guérin), a live attenuated mycobacterium, is currently the only tuberculosis vaccine in use in humans. For more than four decades, intravesical BCG has been the gold standard therapy for high-risk non-muscle-invasive bladder cancer ^16^. Our group recently demonstrated that intravenous BCG delivery reduced tumor growth and prolonged survival in several mouse models of lung tumors. Mechanistically, intravenous BCG promotes NK cell and T cell-mediated antitumor immunity in the lungs, highlighting its potential in the treatment of lung tumors^17^.

In this study, we propose the use of intravenous BCG as a potential treatment for pulmonary metastases induced by bladder cancer cells in murine models. Our data indicated that BCG can prevent the formation of bladder cancer lung metastases, although it was ineffective when administered in the presence of established tumors. We also demonstrated that tumor cells developed specific resistance mechanisms targeting NK cells, with a remarkable contribution of IFN-g response to this process. Characterizing the mechanisms acquired by tumor cells to evade immunotherapy is essential for developing novel antitumor strategies to overcome these barriers.

## RESULTS

### Prophylactic treatment with intravenous BCG impairs tumor development and increases survival

MB49 tumor cells are widely used to generate orthotopic bladder tumors in mice, and their intravenous (i.v.) administration has been shown to induce lung metastasis ^18–22^. The first objective of this study was to optimize the MB49 lung metastasis model in our experimental setup. Our data revealed that all mice inoculated with MB49 cells via the i.v. route developed lung metastases at the time of application of the experimental human endpoint to each mouse. Additionally, we observed a wide variety of macroscopic metastases in different organs, including the subcutaneous tissue, thoracic and abdominal cavities, liver, kidneys, heart, and pancreas (Supplementary Fig. 1a). To further characterize tumor cell dissemination, animals were injected with MB49 cells expressing luciferase, and the endpoint was set at day 15, the time point at which animals began to exhibit symptoms. As shown, tumor cells were primarily located in the lungs in all cases (Supplementary Fig. 1b). Other sites of metastasis included bone, kidneys, and stomach.

We previously demonstrated therapeutic efficacy of intravenous BCG in lung tumor models induced by B16F10 and LLC cells (melanoma and lung cancer respectively)^17^. In the present study, we investigated the efficacy of systemic BCG in the MB49 lung metastasis model. Mice were treated with intravenous BCG prior (BCGpre, prophylactic) or after tumor challenge (BCGpost, therapeutic), as described in the Fig. 1a. Mice inoculated with BCG before tumor challenge showed a marked improvement in survival compared to non-treated mice (53.5 vs 17.5 of median survival). In contrast, administration of BCG after tumor challenge did not provide any therapeutic benefit (Fig 1b).

**Figure 1.**
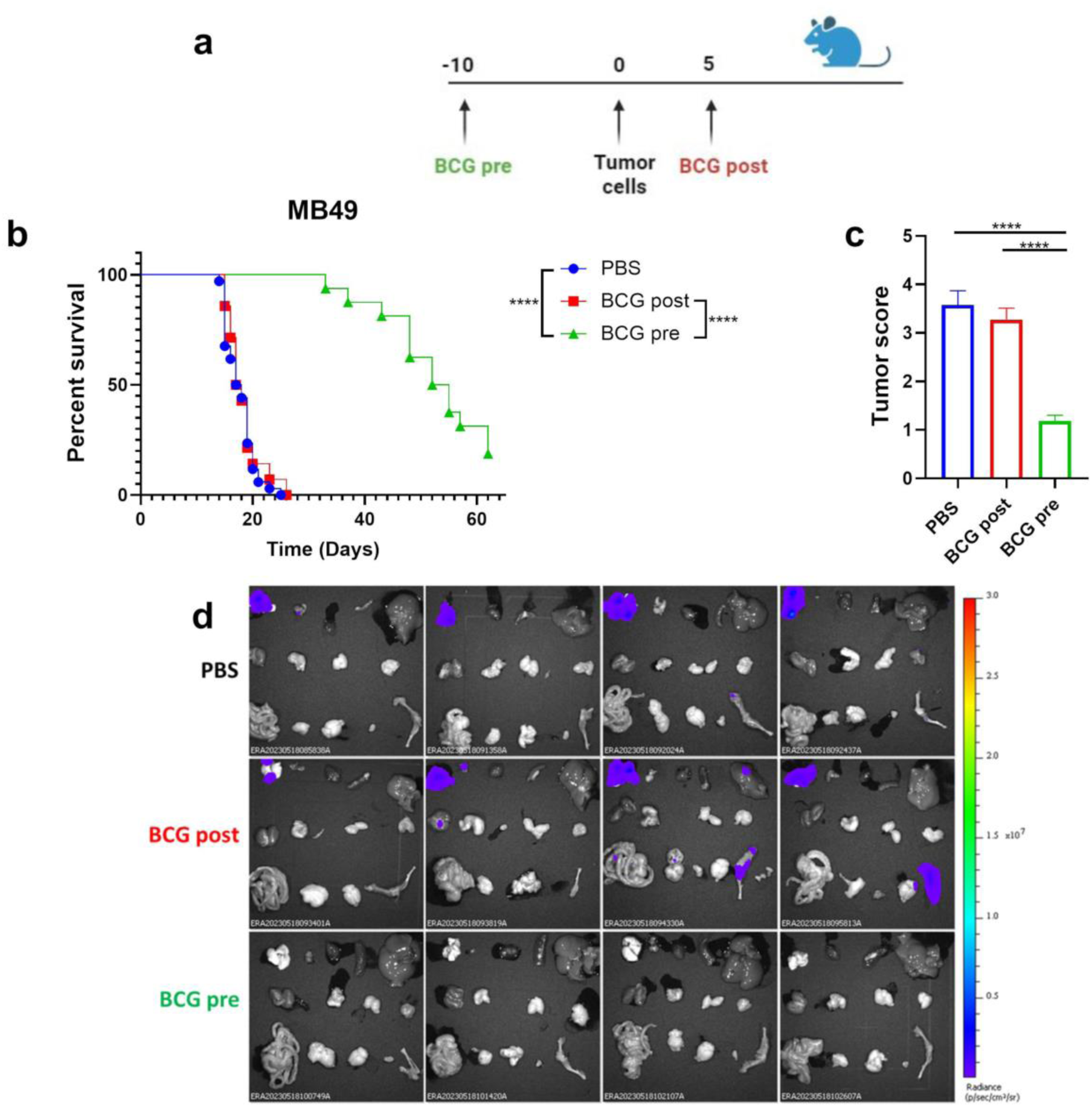
Administration of intravenous BCG prior to tumor challenge increases survival and reduces disease progression. **a.** Schematic diagram showing treatment strategy. **b.** Survival of male mice inoculated with 1×10^5^ MB49 cells intravenously and treated with intravenous BCG after or before tumor challenge. *n*=34 PBS from 5 independent experiments, *n*=17 BCG post, *n*=16 BCG pre, pooled from 4 independent experiments. **** *P* < 0.0001, log-rank (Mantel-Cox) test. **c.** Tumor score of the frequency of number of tumors in distinct anatomical locations observed by visual inspection on day 14 after tumor challenge. *n*=6-12 mice/group pooled from 2 independent experiments. Data depicted as mean ± SEM. **** *P* < 0.0001, one-way ANOVA test. **d.** IVIS imaging for *ex vivo* on day 15 after MB49-ZsGreenLuc i.v. inoculation in male mice treated with intravenous BCG before or after tumor challenge. *n*=4 PBS, *n*=4 BCG post, *n*=4, BCG pre, from 1 independent experiment. First line from left to right: lungs, mediastinal lymph node, heart, spleen and liver. Second line from left to right: kidneys, stomach, genitals and parotid gland. Third line from left to right: intestines, pancreas, brain, bladder, femur and tibia. Radiance: p/sec/cm2/sr. Color scale: Min=3×10^4^, Max=3×10^7^.

To further characterize the efficacy of BCG, necropsies of MB49-tumor-bearing mice were conducted on day 14, and the number of tumors quantified through macroscopic visual examination. Based on these results, we established a tumor score that was used to compare differences between groups. Consistent with the survival results, no differences in tumor number were observed between untreated mice and those treated with BCGpost. However, mice treated with BCGpre displayed a significantly reduced tumor burden compared to the other groups (Fig. 1c). These results suggest that prophylactic intravenous BCG can reduce tumor development and improve survival outcomes. Using MB49-luc cells and the IVIS imaging system, we found that BCGpre completely prevented tumor establishment in the lungs and secondary organs by day 15 post-tumor challenge (Fig. 1d), confirming the strong antitumor effect of BCG as a prophylactic approach.

### Anti-tumor efficacy of BCG relies on IFN-γ, perforin and NK cells

To elucidate the anti-tumor immunological mechanism of prophylactic BCG, the vaccine administered prior to tumor challenge was tested in IFN-γ and Perforin (Perf1) knock-out mice (Fig. 2a). The results showed no difference in survival between treated and control groups in IFN-γ KO mice (Fig. 2b), indicating that IFN-γ is indispensable for BCG’s anti-tumor efficacy. In the case of Perf1 KO mice, survival improvement in the BCG-treated group was minimal but significant (Fig. 2c), suggesting that the BCG-induced anti-tumor response is primarily dependent on perforin-induced cytotoxicity, although other mechanisms might contribute minimally. Altogether, these results emphasize the crucial importance of cytotoxic cells for BCG-mediated tumor efficacy.

**Figure 2.**
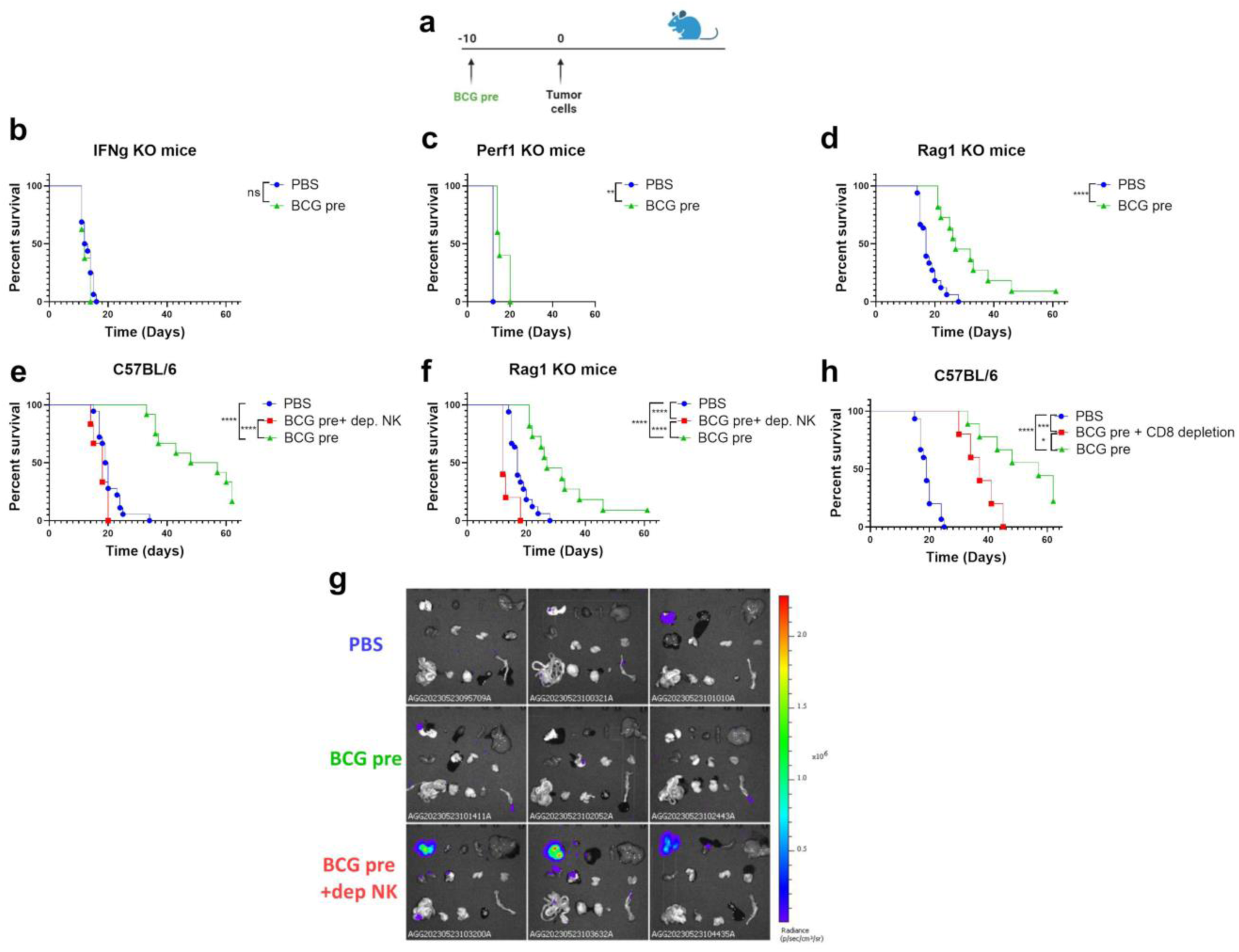
Efficacy of intravenous BCG depends on IFN-γ, perforin and NK cells. **a.** Schematic diagram showing treatment strategy. Survival of males treated with intravenous BCG before inoculation of 1×10^5^ MB49 cells in **b.** IFN-γ KO mice (*n*=4-8 mice/group, pooled from 2 independent experiments), **c.** Perforin KO (*n*=4-5 mice/group, from one experiment).**d.** Rag1 KO mice (*n*=33 PBS, pooled from 6 independent experiments; *n*=10 BCG pre, pooled from 2 independent experiments) and Survival males treated with intravenous BCG before inoculation of 1×10^5^ MB49 cells and treated with αNK1.1 antibody in **e.** WT mice (*n*=18 PBS pooled from 3 independent experiments, *n*=6 BCG pre+dep. NK from 1 experiment, *n*=12 BCG pre pooled from 2 independent experiment) and **f.** Rag1 KO mice (*n*=33 PBS pooled from 6 independent experiments, *n*=5 BCG pre+dep. NK from 1 experiment, *n*=11 BCG pre pooled from 2 independent experiment). **g.** IVIS imaging *ex vivo* on day 6 of male mice after 1×10^5^ MB49-ZsGreenLuc cells inoculation, previously treated with intravenous BCG and NK depleted. n=3 mice/group from one experiment. First line from left to right: lungs, mediastinal lymph node, heart, spleen and liver. Second line from left to right: kidneys, stomach, genitals and parotid gland. Third line from left to right: intestines, pancreas, brain, bladder, and femur and tibia. Radiance: p/sec/cm2/sr. Colour scale: Min=2.36×10^3^, Max=2.29×10^6^**h.** Survival of WT male mice treated with intravenous BCG before inoculation of 1×10^5^ MB49 cells and treated with anti-CD8α. *n*=25 PBS pooled from 5 independent experiments, *n*=5 BCG pre+CD8 depletion from one experiment, *n*=10 BCG pre pooled from 2 independent experiments. ns: not significant, * *P* < 0.05, ** *P* < 0.01, *** *P* < 0.001, **** *P*< 0.0001 log-rank (Mantel-Cox) test.

To assess the differential contribution of CD8+ T lymphocytes and NK cells, we used Rag1 KO mice, which lack T and B cells but retain functional NK cells. Unlike mice lacking IFN-γ or Perf1, Rag1 KO mice treated with BCGpre exhibited a substantial improvement in survival rates compared to untreated mice (Fig. 2d). These results suggested a strong contribution of innate cytotoxic cells to BCG preventive efficacy.

To confirm the involvement of NK cells, we depleted this population using specific neutralizing antibodies. Results in both wild-type and Rag1 KO mice revealed that NK cell depletion abrogated the anti-tumor effect of BCG (Fig. 2e, f), a finding confirmed by IVIS imaging at day 6 post-tumor cell inoculation (Fig. 2g). Additionally, depletion of CD8+ T cells in wild-type mice had a negligible impact on the survival of BCGpre-treated mice compared to non-depleted animals (Fig. 2h), which aligned with our findings in Rag1 KO mice. These data demonstrate the pivotal role of NK cells, rather than CD8+ T lymphocytes, in the BCG-mediated response against MB49 cell lung metastasis.

### *In vitro* incubation of MB49 cells with IFN-g increases tumor immunotherapy resistance

Our results indicated a strong anti-tumor prophylactic effect of BCG, primarily through NK cell activation. However, the lack of efficacy when BCG was administered after tumor establishment suggests that the tumor had already triggered mechanisms to evade BCG-activated immunity. In recent years, growing evidence has highlighted the crucial role of interferon signaling in promoting tumor resistance to immunotherapy, primarily through the upregulation of checkpoint molecules such as PD-L1^23^. Given that BCG is a potent inducer of IFN-γ, we hypothesized that this cytokine might mediate the resistance of MB49 cells to BCG. ^23^

To investigate this, we firstly studied the responsiveness of MB49 cells to IFN-γ by incubating the cells with the cytokine and analyzing the levels of MHC-I, PD-L1, and MHC-II, all of which are known to be upregulated by IFN-γ. Our data revealed a high basal expression of MHC-I and minimal or no expression of PD-L1 and MHC-II, respectively (Fig. 3a-c). Incubation with IFN-γ led to a substantial increase in surface expression of all three markers, confirming that MB49 cells are sensitive to IFN-γ.

**Figure 3.**
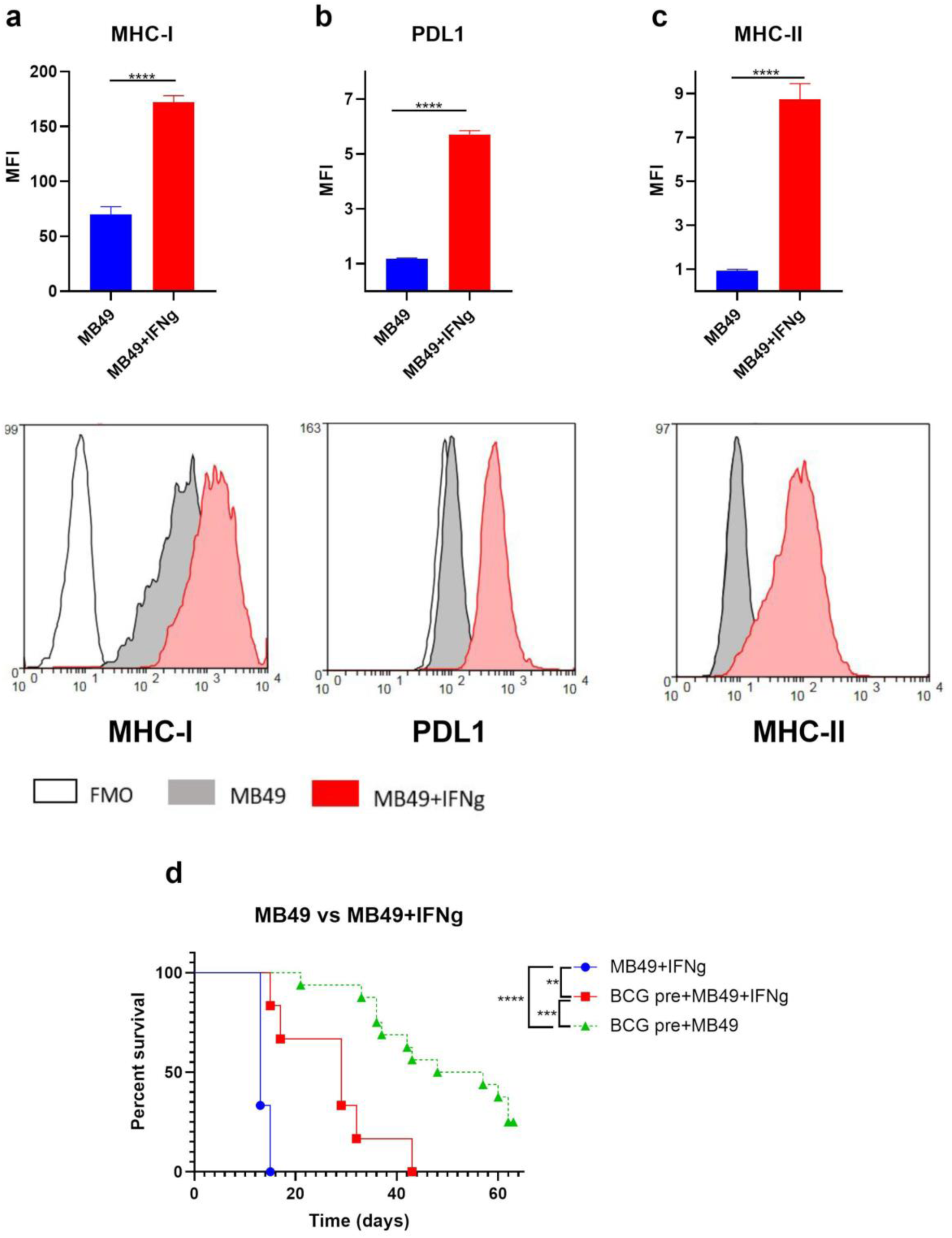
Stimulation of MB49 cells with IFN-γ reduces the efficacy of prophylactic intravenous BCG. Tumor cells were incubated with IFN-γ for 24 hours and IFN-γ signature was analyzed by flow cytometry. Expression and histograms of MB49 expression of (**a**) MHC-I, (**b**) PDL1 and (**c**) MHC-II. MFI was expressed as MFI of each replicate divided to FMO. **** *P* < 0.0001, unpaired t-test. **g.** Survival of MB49 stimulated with IFN-γ *in vitro* tumor-bearing male mice treated with intravenous BCG after tumor challenge. *n*=6 mice/group for MB49+IFN-γ and BCG pre+MB49+IFN-γ, from one experiment. *n*=16 mice/group for BCG pre+MB49, pooled from 3 independent experiments. ** *P* < 0.01, *** *P* < 0.001, **** *P* < 0.0001, log-rank (Mantel-Cox) test.

To assess the potential role of IFN-γ in mediating MB49 resistance to BCG, we incubated MB49 cells with IFN-γ for 24 hours and subsequently inoculated them into mice previously treated with intravenous BCG. Our data revealed a significant decrease in survival in mice inoculated with IFN-γ-treated MB49 cells compared to those inoculated with naïve tumor cells (Fig. 3d). These findings suggest that IFN-γ plays a role in editing MB49 tumor cells to develop resistance to immunotherapy.

### IFNg signalling depletion partially restores BCG therapeutic efficacy

Based on these results, we hypothesized that IFN-γ might mediate MB49 cell resistance in the therapeutic context. To address this, we disrupted the JAK1 gene in MB49 cells (MB49 JAK1 KO) using CRISPR/Cas9, effectively abolishing the primary IFN-γ signaling pathway. The lack of upregulation of MHC-I, PD-L1, and MHC-II following incubation with IFN-γ for 24 hours (Fig. 4a) confirmed that MB49 JAK1 KO cells were unresponsive to IFN-γ. We then inoculated these cells into mice and subsequently treated them with either PBS or BCG on day 5. Unlike the results observed with wild-type MB49 cells, BCG-treated mice inoculated with MB49 JAK1 KO cells exhibited significantly higher survival compared to untreated mice (Fig. 4b), suggesting the relevance of IFN-γ in promoting immune evasion functions that support tumorigenesis and subvert anti-tumor responses.

**Figure 4.**
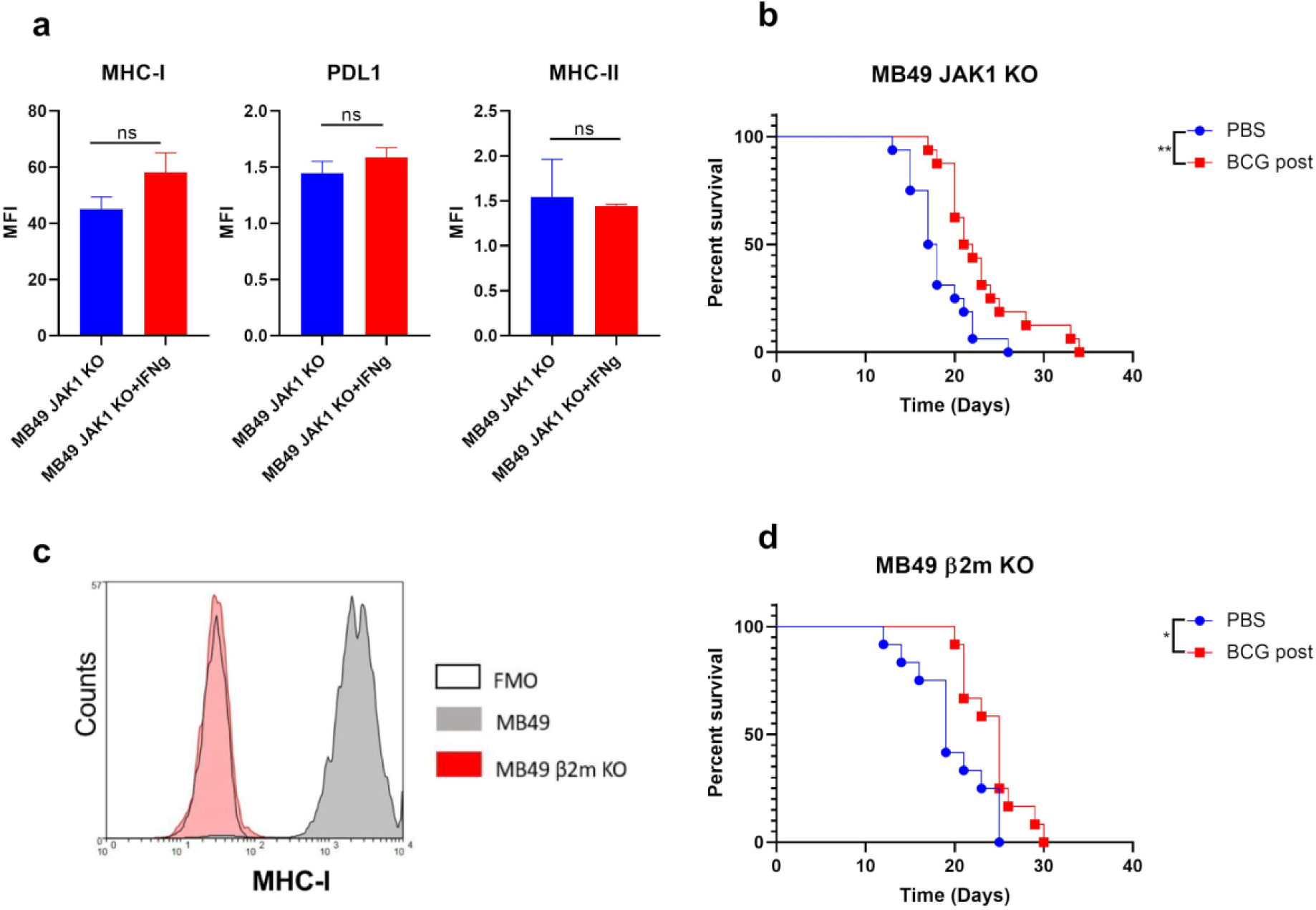
Lacking of MHC-I and disruption of IFN-γ signaling pathway leads to a partial recovery of intravenous BCG efficacy. **a.** Expression of MHC-I, PDL1 and MHC-II, analysed by flow cytometry, of MB49 and MB49 JAK1 KO cells after incubation with IFN-γ for 24 hours.**b.** Survival of MB49 JAK KO-tumor-bearing male mice treated with intravenous BCG after tumor challenge. *n*=12 mice/group, pooled from 2 independent experiments. ** *P* < 0.01, log-rank (Mantel-Cox) test. **c.** Histogram of MHC-I expression of MB49 and MB49 β2m KO. **d.** Survival of MB49 β2m KO-tumor-bearing male mice treated with intravenous BCG after tumor challenge. *n*=12 mice/group, pooled from 2 independent experiments. * *P* < 0.05, log-rank (Mantel-Cox) test.

Given our findings on the importance of NK cells in BCG-mediated immunosurveillance and the high surface expression of MHC-I by MB49 cells, we hypothesized that the elevated MHC-I levels inhibited the cytotoxic function of BCG-activated NK cells, thereby contributing to treatment inefficacy. To test this, we disrupted the β2-microglobulin gene in MB49 cells (MB49 β2m KO), which abrogated surface MHC-I expression (Fig. 4c). *In vivo* results showed a significant improvement in the efficacy of intravenous BCG in mice challenged with MB49 β2m KO cells (Fig. 4d), suggesting that MHC-I expression contributes to MB49 lung tumor immune evasion.

## DISCUSSION

While immunotherapy has revolutionized cancer treatment over the past decade, its widespread use has also led to an increase in immunotherapy resistance. In the case of lung tumors, immunotherapy resistance affects until 60% of patients treated^24^. Understanding how immunotherapy itself edits tumor cells to acquire additional resistance mechanisms is vital for the development of more effective therapies ^25^. In the current study, we identified two contrasting scenarios: one in which immunotherapy was highly effective (when administered before tumor challenge) and another where it failed (when administered after tumor challenge). The comparison between these two situations could allow to identify effective tumor immunosurveillance mechanisms, as well as tumor immune evasion strategies.

We selected intravenous BCG as immunotherapy, a treatment that we have previously reported to have therapeutic efficacy in other lung tumor models, such as those induced by B16-F10 melanoma cells, through a mechanism highly dependent on NK cell activation ^17^. This finding is consistent with the present study, where we observed, in a lung metastasis model induced by bladder cancer cells, a crucial role for NK cells in preventing lung metastasis, as evidenced by depletion studies. The data indicated that the prophylactic antitumor capacity of BCG relied heavily on NK cells, with a limited contribution from CD8+ T lymphocytes. Notably, the prophylactic efficacy of BCG was not exclusive to the MB49 model, as we previously observed similar results in mice challenged with B16-F10 cells^17^. Given the importance of circulating NK cells in eliminating tumor cells ^4^, we speculate that BCG-activated NK cells may eliminate MB49 cells before they reach the lungs, which would explain the high prophylactic efficacy observed. In contrast, once lung tumors are established, it is likely that tumor cells have already deployed escape mechanisms, rendering the BCG-induced immune response ineffective.

A relevant question that arises from comparing this study with our previous one is why BCG does not work against MB49 cells, while it is effective in treating tumors bearing B16-F10 cells. One important consideration is that these differences between cell lines could be instrumental in identifying immunotherapy escape mechanisms. For instance, a key difference between MB49 and B16-F10 cells is the basal expression of MHC-I, which is very high in MB49 cells, whereas B16-F10 cells barely express it (although it is inducible by IFN-γ in both cases). We hypothesize that the high level of MHC-I on the surface of MB49 tumor cells may impair the activity of NK cells when they encounter the tumor. Indeed, our data with MB49 β2m KO cells (which are unable to express MHC-I) showed partial recovery of BCG’s therapeutic efficacy, suggesting that inhibition of NK cells through MHC-I expression could be a mechanism of immune escape for MB49 cells.

Our results also suggest that tumor evasion mechanisms are not fully intrinsic to MB49 cells, but are at least partially acquired *in vivo*. This is indicated by two key findings: First, BCG is highly effective when administered before MB49 cell inoculation, meaning tumor cells are rapidly eliminated when the antitumor response has been previously boosted by BCG, regardless of MHC-I basal levels. Second, MB49 cells preconditioned with IFN-γ exhibited resistance to BCG-induced antitumor immunity. This finding highlights the importance of IFN-γ in enabling tumor cells to escape immunotherapy, a mechanism that has been previously reported ^26^. Recent clinical studies have shown that late inhibition of IFN-γ pathways with specific JAK inhibitors can help overcome tumor resistance to immune checkpoint inhibitor-based immunotherapy ^27,28^. In the context of our studies, we demonstrated that MB49 cells deficient in JAK1 (and therefore unresponsive to IFN-γ) were less resistant to therapeutic BCG, corroborating the importance of IFN-γ in mediating resistance to BCG’s antitumor response. This correlates with the strong induction of checkpoint molecules such as MHC-I and PD-L1 observed in MB49 cells incubated *ex vivo* with IFN-γ. IFN-γ-mediated expression of checkpoint molecules has been linked to immunotherapy resistance in various studies ^23^.

Our findings may have important implications for designing strategies aimed at preventing or managing lung metastasis from bladder cancer or other primary tumors. Neoadjuvant immunotherapy has shown promising efficacy in several cancer types, such as lung and hepatocellular carcinoma, by preventing recurrences and metastasis and extending patient survival ^29,30^. Our observation of BCG’s prophylactic efficacy in preventing lung metastasis suggests that intravenous BCG could be a promising neoadjuvant approach to prevent lung metastasis following the resection of primary tumors.

## METHODS

### Ethics statement

The use and care of animals for experimental work were performed in agreement with the Spanish Policy for Animal Protection RD53/2013 and the European Union Directive 2010/63 for the protection of animals used for experimental and other scientific purposes. Experimental procedures were approved by Ethics Committee for Animal Experiments of University of Zaragoza (PI46/18, PI33/15 and PI50/14).

### Mouse strains

C57BL/6JR mice were purchased to Janvier Biolabs or bred in the facilities of the Centro de Investigaciones Biomédicas de Aragón (CIBA). Mouse strains deficient in IFN-γ, Rag1, Batf3 and Perforin were bred in the facilities of the Centro de Investigaciones Biomédicas de Aragón (CIBA). Male and female mice aged 8 to 12 weeks were used for the experiments. Mice were acclimatized during one-two weeks before the experiments started. Food and water were provided ab libitum. Room temperature was 20-24°C, humidity 50-70% and the light intensity 60 lux with the light-dark cycle of 12 hours.

### Tumor cell lines

The MB49 cell line is a urothelial carcinoma line originally obtained by exposing bladder epithelial cells obtained from a C57BL/Icrf-a’ mouse to the carcinogen 7,12-dimethylbenz[a]anthracene (DMBA) for 24 hours. MB49 cells used in our laboratory were purchased from ATCC. MB49 cell line authenticity was checked by Short Tandem Repeat (STR) analysis (Multiplexion). MB49 expressing the fluorescent protein ZsGreen and the reporter luciferase (MB49-ZsGreenLuc) were made in our laboratory by transfection with a lentivirus encoding ZsGreen and firefly luciferase Luc2P (pHIV-Luc2-ZsGreen, from Addgene) and sorted based on high ZsGreen expression. For the generation of MB49-*β2m^-/-^* and MB49-*JAK1^-/-^*, parental cells were transfected with CRISPR-Cas9 plasmids targeting the β2-microglobulin gene and Janus kinase 1 (purchased from SantaCruz Biotechnology) and cells were selected with puromycin. CRISPR plasmids containing three different sgRNAs targeting the β2-microglobulin gene (in case of lacking MHC-I expression) or JAK1 gene, a puromycin resistance gene and RFP for selection of transfected populations. Then transfected cells were sorted based on lack of MHC-I expression after staining with an APC-conjugated H2K^b^/D^d^ antibody, and sorter again, if necessary, in case of MB49-*β2m^-/-^.* In case of MB49 JAK1 KO, the first sorting was based on RFP expression due to the inability of sorting based on an intracellular protein like JAK1. The second sorting was based on lack of PD-L1 expression after staining with an PE-conjugated PD-L1 antibody. In this case, MB49 transfected cells were stimulated with IFN-γ during 24 hours, then, stained with PE-conjugated PD-L1 (BD) antibody and sorted based on lack of PD-L1 expression. All cell lines were cultured in DMEM (Gibco) supplemented with 10% (v/v) heat inactivated fetal bovine serum (FBS, Gibco), 1% (v/v) L-glutamax (Sigma), 100 U/ml penicillin (Sigma), 100 U/ml streptomycin (Sigma) and 10 μg/ml ciprofloxacin (Santa Cruz Biotechnology). Cell lines were routinely passaged by addition of trypsin/EDTA (Sigma) and were always used with less than 10 passages from thawing. Cell lines were stored frozen in liquid nitrogen at a concentration of 2×10^6^ cells/ml in heat inactivated FBS (Gibco) supplemented with 10% dimethyl sulfoxide (Sigma-Aldrich). Mycoplasma contamination was routinely checked in cell culture supernatants.

### Experimental murine tumor models

For tumor cell inoculation, cells were cultured as described, detached with trypsin/EDTA, counted and resuspended at the desired concentrations in serum-free DMEM. For the induction of metastasis in lungs, 1×10^5^ MB49 or MB49-derived tumor cells were inoculated into the tail vein of mice in a volume of 200 μl in serum-free DMEM.

For survival experiments, mice were euthanized by cervical dislocation upon reaching the humane endpoint. Mice were monitored twice a week.

For the IVIS and tumor score experiments, mice were euthanized at a predetermined point (between days 14 and 15) when they began to show symptoms. For the tumor score experiments, necropsies were performed, and tumor progression was assessed by counting the number of tumors in the lungs and the number of metastases in different organs. A tumor score was established based on the number of tumors counted by visual inspection: 1 point, 0-25 tumors; 2 points, 25-50 tumors; 3 points, 50-75 tumors; and 4 points, 75 tumors or more.

### Bacterial strains

The BCG Pasteur reference strain 1173P2 used in this study, was given from Roland Brosch (Institut Pasteur, France). BCG strains were grown at 37°C in Middlebrook 7H9 broth (BD Difco) supplemented with 0.05% Tween 80 (Sigma) and 10% Middlebrook albumin dextrose catalase enrichment (ADC, BD Biosciences), or in solid Middlebrook 7H10 agar (BD Difco) supplemented with 10% ADC (BD Biosciences). Mycobacteria were grown to mid-log phase in liquid 7H9 supplemented broth, and culture were centrifugated and resuspended in PBS with 0.05% Tween 80. Bacterial suspensions were kept for 10 minutes at room temperature to allow clump sedimentation, and the resulting supernatants were centrifugated to remove additional clumps at 800g for 10 minutes. Finally, glycerol was added to supernatants to achieve a final concentration of 5%. Aliquots were stored at - 80°C. One week after freezing the cultures, a vial of the batch was thawed, plated on solid Middlebrook 7H10 agar and 3 weeks later colonies were counted to determine CFU concentration of each batch. All experiments were performed with bacteria from quantified stocks kept at −80°C. Bacteria from frozen quantified stocks were resuspended in PBS. Part of the inoculum used for treatment was plated in solid medium for CFU determination and quality control. Mice were inoculated with 10^6^ CFUs of bacterial suspensions into the tail vein in a volume of 200 μl.

### Antibody-based cell depletion and treatments

For CD8+ T cell and NK cell depletion, mice were injected intraperitoneally with 100 μg of anti-CD8α (clone 2.43, BioXCell) or 100 μg of anti-NK 1.1 (clone PK136, BioXCell) twice a week, initiating the procedure one day before tumor challenge.

### Image studies

To study the tumor progression and dissemination in vivo, the follow-up of tumor-bearing mice was carried out using IVIS Lumina LT In Vivo Imaging System in the facilities of Centro de Investigaciones Biomédicas de Aragón (CIBA). For these studies, mice were inoculated with MB49-ZsGreenLuc intravenously and the follow-up was carried out at different time points depending on the experiment. Mice were administered with 150 ug/g of D-luciferin potassium salt (Deltaclon) intraperitoneally, followed by image acquisition 8 minutes later. Organs were distributed as indicated in Supplementary Figure 2

### IFN-γ signature assay

1×104 tumor cells were seeded in flat 96-well plates and incubated during 24 hours. Then, 50 ng/ml of IFN-γ were added and incubated during 24 or 48 hours depending on the experiment conditions. After, tumor cells were stained using APC-conjugated-MHC-I (Miltenyi), Vioblue-conjugated-MHC-II (Miltenyi) and PE-conjugated-PDL1 (BD) antibodies and analyzed by flow cytometry.

### Flow cytometry

Fluorescence minus one (FMO) stained controls were used for the discrimination of negative and positive populations and for the correction of fluorescence spill-over of the different fluorochromes. Data concerning optimization and compensation of the different flow cytometry antibody panels used in this study is not shown although it was performed for each panel. Single cell suspensions were plated in U-bottom 96-well plates. Firstly, single cell suspensions were incubated with mouse Fc receptor blocking reagent (Miltenyi) for 20 min at 4°C in PBS 2% FBS. Then, surface staining was performed for 20 minutes at 4°C using the following antibodies anti-H2K^b^/D^b^-APC (clone REA932, Miltenyi), anti-MHC-II-Vioblue (clone REA813, Miltenyi) and anti-PD-L1-PE (clone MIH5, BD). Cells were acquired using a Gallios Flow cytometer (Beckman Coulter). Flow cytometry results were analyzed using Weasel software (3.0.2 version).

### Data Analysis

Mice were distributed in groups of at least 3-6 animals per cage prior to experimental procedures. Results were not blinded for analysis and randomization was not applicable to these studies. GraphPad Prism software (version 8.0.2) was used for graphical representation and statistical analysis. Statistical tests used in each experiment are indicated in the figure legends. To compare two group means, unpaired Student’s t-test was used. To compare means between more than two groups, unpaired one-way ANOVA was used. For survival analysis, log-rank Mantel-Cox test was used. P values < 0,05, were considered significant: * p < 0.05, ** p < 0.01, *** p < 0.001, **** p < 0.0001. Data shown in graphs depicts mean +/- standard error of the mean (SEM).

### Data availability

Materials are available upon request to the authors.

## ACKNOWLEDGMENTS

This work was supported by MCIN/AEI/10.13039/501100011033 [grants number RTI2018-097625-B-I00 and PID2022-138624OB-I00], Gobierno de Aragón [grant number LMP50_21] and Asociación Española Contra el Cáncer (AECC) [grant number IDEAS211042AGUI]. NA was the principal investigator of all these grants. This research was supported by CIBER-Consorcio Centro de Investigación Biomédica en Red-(Groups CB06/06/0020), Instituto de Salud Carlos III, Ministerio de Ciencia e Innovación and Unión Europea-European Regional Development Fund. CG has a pre-doctoral fellowship from Gobierno de Aragón. The funders had no role in study design, data collection and analysis, decision to publish or preparation of the manuscript. Authors acknowledge the Scientific and Technical Services from Instituto Aragonés de Ciencias de la Salud (IACS) and Universidad de Zaragoza.

## AUTHOR CONTRIBUTIONS

C.G., C.M., and N.A. designed the experiments. C.G., E.M., S.U., A.B.G., and M.A-V. performed the experiments. C.G. and N.A wrote the manuscript. N.A. supervised the study.

