## Supplementary material for "NK cells mediate preventive efficacy of intravenous BCG against lung metastasis in mice"

**SUPPLEMENTARY INFORMATION**

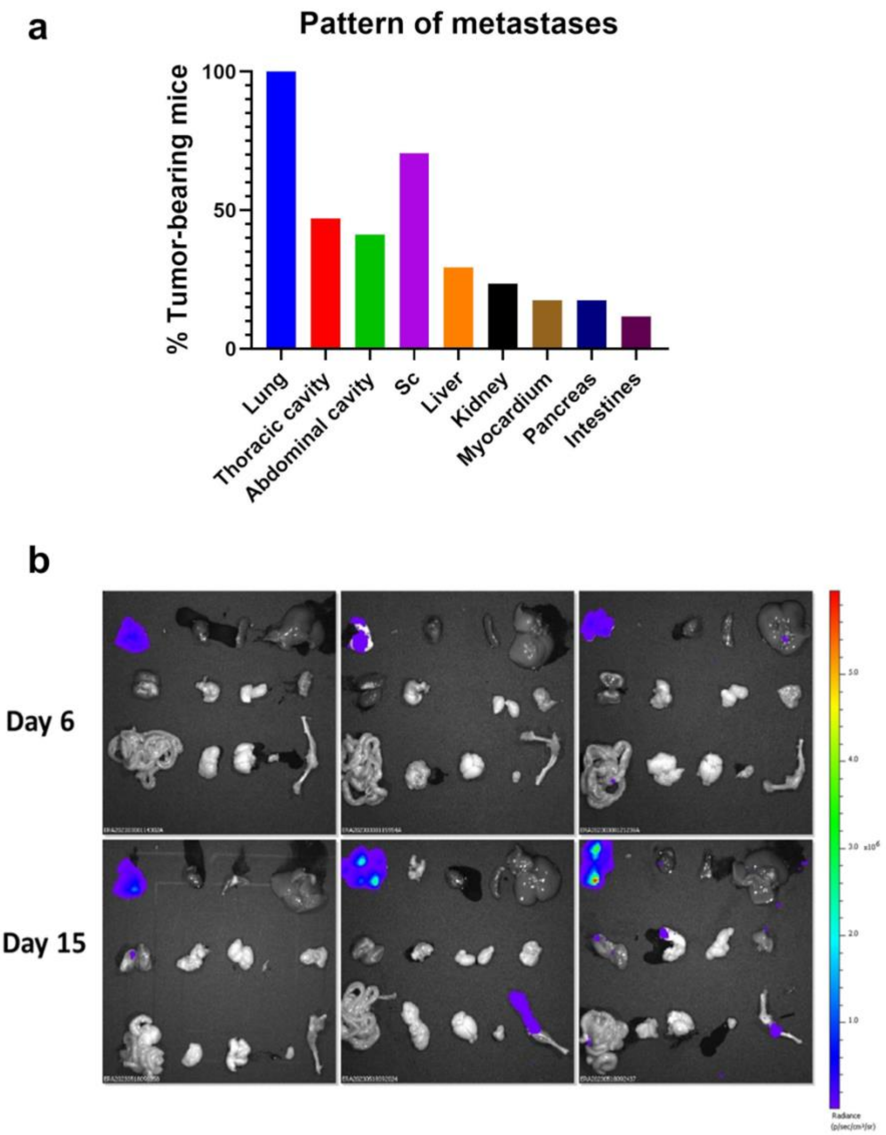


**Supplementary Figure 1. Intravenous administration of MB49 cells causes metastases into mice. a.** Frequency of metastases found in distinct anatomical locations observed by visual inspection in MB49 tumor-bearing male mice. Mice were euthanized when humane endpoint was reached. *n*=17 mice, pooled from 3 independent experiments**. b.** IVIS imaging for *ex vivo* on day 6 and 15 after MB49-ZsGreenLuc i.v. inoculation in male mice. First line from left to right: lungs, mediastinal lymph node, heart, spleen and liver. Second line from left to right: kidneys, stomach, genitals and parotid gland. Third line from left to right: intestines, pancreas, brain, bladder, and femur and tibia. Radiance: p/sec/cm2/sr. Color scale: Min=9.76x10$^{3}$, Max=5.96x10$^{6}$.

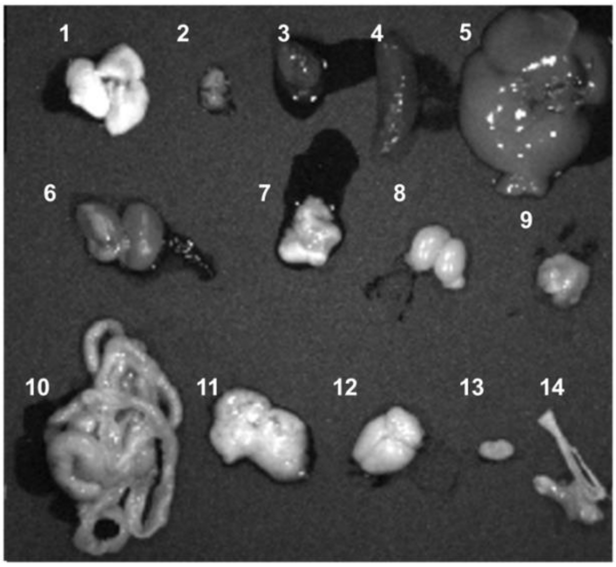


**Supplementary Figure 2. Distribution of organs for IVIS image studies**. 1: lungs, 2. mediastinal lymph nodes, 3: heart, 4: spleen, 5: liver, 6: kidneys, 7: stomach, 8: genitals, 9: parotid gland, 10 intestines, 11: pancreas, 12: brain, 13: bladder, 14: femur and tibia.
